# CLIMBS: assessing carbohydrate-protein interactions through a graph neural network classifier using synthetic negative data

**DOI:** 10.1101/2025.02.27.640667

**Authors:** Yijie Luo, Fabio Parmeggiani

**Affiliations:** School of Biochemistry, University of Bristol; University Walk, Bristol BS8 1TD, UK; School of Chemistry, University of Bristol; Cantock’s Close, Bristol BS8 1TS, UK; School of Pharmacy and Pharmaceutical Sciences, Cardiff University, Redwood Building, King Edward VII Ave, Cardiff, CF10 3NB, UK

## Abstract

Carbohydrate-protein interactions are essential for biological processes, such as cellular signaling and metabolism, and represent a large pool of untapped targets for diagnostics and therapeutics. However, current design and prediction methods fail to accurately evaluate the affinity and specificity of proteins for carbohydrates such as glucose and galactose. Here, we describe a machine learning classifier, named CLIMBS, as a novel scoring method for protein-carbohydrate interactions and train it on crystal structures and synthetic data from unsuccessfully designed binders, to effectively assess if carbohydrate-protein complexes represent realistic, native like structures. Compared to other methods, CLIMBS has outstanding accuracy, excellent carbohydrate specificity, sub-second runtime per sample, minimal bias towards either negative or positive samples, and can be employed to improve selection of successful docking and design models of carbohydrate-protein complexes.

## II. Introduction

Carbohydrate recognition by proteins is a fundamental step in a number of biological processes, from glycolysis(*1*) to assembly of specific glycosylation patterns and formation of polysaccharide materials(*2*), either for energy storage, as for starch(*3*), or for structural use, like cellulose(*4*). On the other hand, large groups of lectins act as receptors for glycans and poly-saccharides and play key roles in cellular signalling and adhesion (*5*). Identification of receptors and binding partners, and specificity predictions have become fundamental in understanding interactions between the glycome and the proteome. Although high throughput (*6*) and glycan specificity computational models have been developed(*7, 8*), most of the molecular details for specific interactions remain routinely out of experimental grasp, and addressed through docking and modelling of small molecule-protein interactions(*9, 10*).

Accurately assessing protein-carbohydrate interactions remains challenging for energy-based modelling methods. Even established scoring functions, such as AMBER (*11*), Rosetta (*12*) and AutoDock4 (*13*) can fail to capture weak but key interaction such as CH-π(*14*) when considering the whole complex or interface. Alternatively, machine learning methods have been applied to docking and evaluation of protein-small molecule complexes. GNINA (*15*) and Sfcnn (*16*) use convolutional neural networks (CNN) and present complex structures in 3D-grid voxels. CSM-lig (*17*) and DeepDOCK (*18*) use graph neural networks (GNN), presenting complex structures as graphs with additional physicochemical characteristics as features. Generative models such as DIFFDOCK (*19*) and AlphaFold3 (*20*) have intrinsic confidence models to rank and assess the quality of the generated structures. Some methods have introduced specific biases to explicitly address protein-carbohydrate interactions, for example the GLYCAM06 energy function (*21*) and CSM-carbohydrate model (*22*).

The development of machine learning methods targeted to protein-carbohydrate interactions has the potential to improve assessment of native-like interface in docking(*23*) and design (*24*). However, for protein carbohydrate complexes, there is no available large dataset which unsupervised learning methods can rely on. When looking at available structural information, among 211,432 protein structures in the protein data bank (PDB) (*25*), only 12.33% (26,705 structures) of them contained carbohydrates as of February 2024. And if we consider redundancy, structures with resolution lower than 3 Å and only structures where the carbohydrate form part of a binding interface, the number drops to 3.84% (8,119 structures). In contrast, supervised learning methods, such as classifiers, require much smaller datasets to achieve reasonable performances(*26*), although appropriately labelled. While selected protein carbohydrate-complexes in the PDB can provide positive samples (“binding”), negative samples (or “not-binding”) for complexes, where the carbohydrate is positioned unfavorably in a binding pocket, can only be computationally generated. Interestingly, reasonable non binders can be easily generated, since even state of the art protein design methods have achieved success rates of only 0.67% (*27*) for small molecule binders. This implies that a large synthetic dataset of designed binders for target carbohydrates could provide realistic and challenging negative samples for training supervised learning methods. Building on this hypothesis, we have developed a graph neural network (GNN)-based model: CLIMBS, **cl**ass**i**fier of **m**inimum **b**inding site for **s**ugars (Figure 1). We trained it using positive experimental data and synthetic negative data and then compared its performance in discriminating native and computationally generated complexes with those of established scoring methods. We then explored the potential for retraining on new types of sugars and discriminate complexes with untrained sugars.

**Figure 1.**
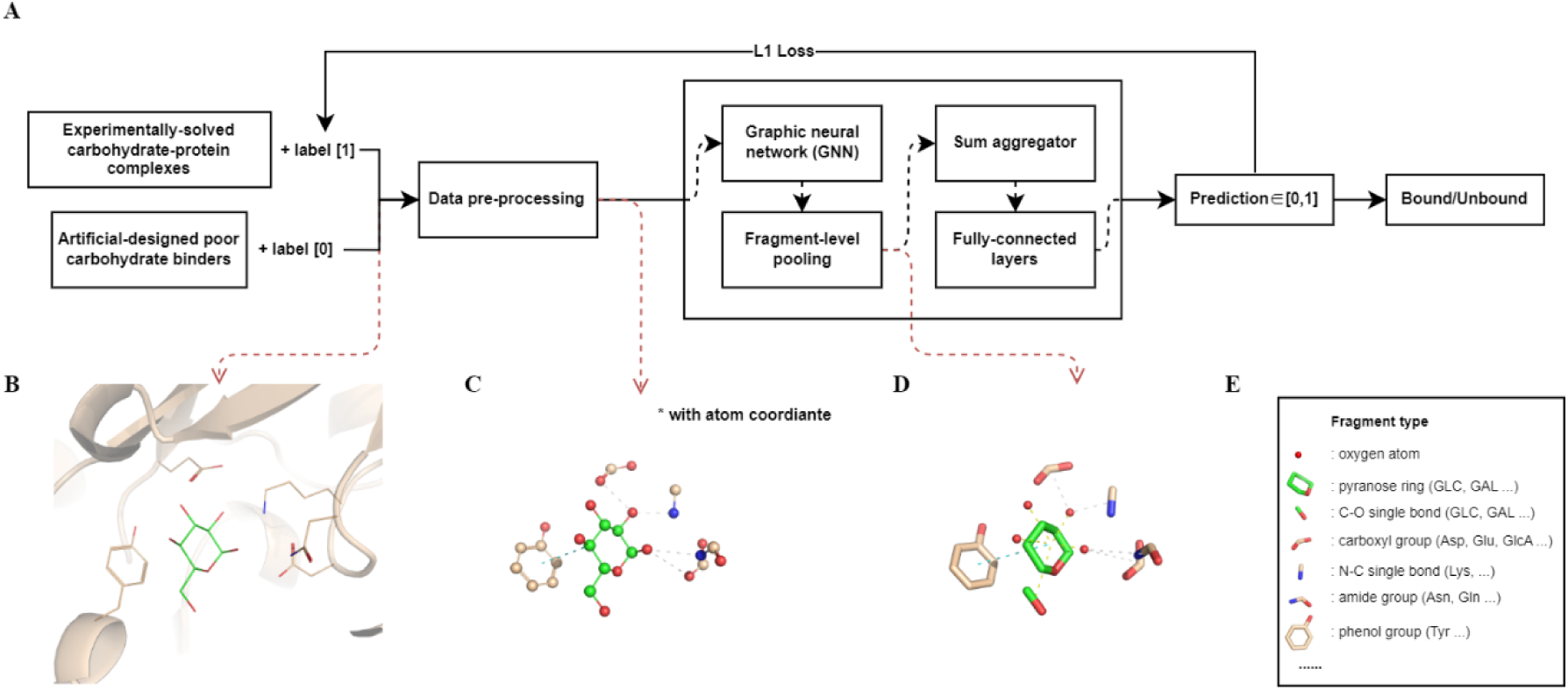
Flowchart of CLIMBS. (A) Overview of CLIMBS. (B) PDB file as input structure data of the model. (C) Structure data becomes an atom-level graph with atom coordinates for each node after data pre-processing. (D) In CLIMBS, structure data becomes a fragment-level graph after a pooling layer. (E) Legend of fragment types shown.

## III. Results

### Overview of CLIMBS

CLIMBS was developed as a graph neural network classifier to validate atomistic models of carbohydrate-protein complex. CLIMBS takes carbohydrate-protein complex structures as inputs, and it outputs 1 or 0 indicating predictions as “binding” or “non-binding”. The binding interface, rather than the whole complex, is used as the model input to focus on binding site geometry and reduce run time. Atom-level geometric information is used in the form of graphs to capture details of interactions. Due to the structural similarity between carbohydrates, fragment-level information (including groups of neighbouring atoms, such as hydroxyl groups or carbon rings, see Figure 1) is processed within a pooling layer, facilitating learning across various types of sugars with shared features (*28, 29*).

A structure library(*30*) was built to include 19 different carbohydrates from experimentally determined carbohydrate-protein complexes that were used as positive samples, while suboptimal computationally-designed carbohydrate-protein complexes for the same carbohydrates were introduced as negative samples (see details in Methods).

The library (see Supplementary Table 1) mirrors the Protein Data Bank, with seven mono-saccharide types (glucose, glucosamine, galactose, fucose, mannose, sialic acid and fructose) accounting for 79% of all complexes.

CLIMBS predicts binding between carbohydrate and proteins with an overall accuracy of 89.42%. The model was trained on 894 positive and 869 negative samples (*db_w1*, see Supplementary Data 3a) with 295 epochs. The binding predictions of different types of sugar-protein complexes were tested on a separate set of complexes containing either carbohydrates used during training or unseen sugars (see table 1). A minimal accuracy of 72.22% was observed for di-saccharides constructed by N-acetyl-β-D-galactosamine and β-D-glucuronic acid (not used in training), and a maximal accuracy of 100% for sucrose. Carbohydrates less represented in the library, such as ribose and arabinose, still reach accuracies similar to those of the most abundant ones.

**Table 1.** Accuracy of CLIMBS tested by different sugar-protein complexes. 14 types of sugar-protein complexes were used for training and 5 types were not included in training. For trained sugar types, tested sample number is the size of the test set after splitting (1/5 of the whole dataset). For untrained sugar types, the sample number is the size of the whole dataset.

| **Sugar type in complex** | **Accuracy**<br>**(295 epochs)** | **Tested**<br>**Sample**<br>**number** |
| --- | --- | --- |
| $\alpha/\beta$-D-glucose | **91.79%** | **390** |
| N-acetyl-$\alpha/\beta$-D-glucosamine | **79.80%** | **99** |
| β-D-galactose | **90.28%** | **72** |
| maltose, GLC-(1,4)-βGlc | **94.44%** | **72** |
| α-D-mannose | **93.75%** | **64** |
| N-acetyl-$\alpha/\beta$-D-galactosamine | **87.88%** | **33** |
| N-acetyl-α-neuraminic acid | **86.21%** | **29** |
| α-L-fucose | **88.00%** | **25** |
| β-D-fructose | **88.00%** | **25** |
| GlcNAc-(1,4)-βGlcA & GlcA-(1,3)-βGlcNAc (repeat unit of heparan sulphate, untrained) | **72.22%** | **18** |
| α-D-ribose (untrained) | **91.67%** | **12** |
| sucrose, GLC-(1,2)- βFru | **100.00%** | **12** |
| 6-O-phosphono-$\alpha/\beta$-D-glucose | **81.82%** | **11** |
| N-acetyl-4-O-sulfo-β-D-galactosamine (untrained) | **72.73%** | **11** |
| N,O6-disulfo-glucosamine | **90.00%** | **10** |
| α-L-arabinose (untrained) | **88.89%** | **9** |
| α-D-xylose | **88.89%** | **9** |
| β-D-glucuronic acid | **87.50%** | **8** |
| Repeated region of unmodified heparan sulphate (constructed by 3 GlcA and 3 GlcNAc, untrained) | **87.50%** | **8** |

A robustness check (see Methods) revealed that both CH-π and polar interactions contribute to classification. Moreover, the trained CLIMBS model can be retrained to include new sugar types whilst maintaining excellent performance when adding only tens of new samples in the training set.

### Performance comparison with other methods

CLIMBS has shown an outstanding performance in predicting sugar binding when compared to other scoring methods, according to runtime and four model performance metrics (accuracy, precision, specificity and sensitivity) (Figure 2). A group of complexes in test set were selected as evaluation set for performance comparison (*db_eval*, see Supplementary Data 3j). We chose score functions from energy-based methods (Rosetta Energy Function REF (*12*), Autodock score function (*13*), HADDOCK score function (*31*)) and a confidence model from a machine learning approach (DIFFDOCK (*19*)). Each method was calibrated with a threshold value to provide the highest accuracy (see Supplementary Data 5).

**Figure 2.**
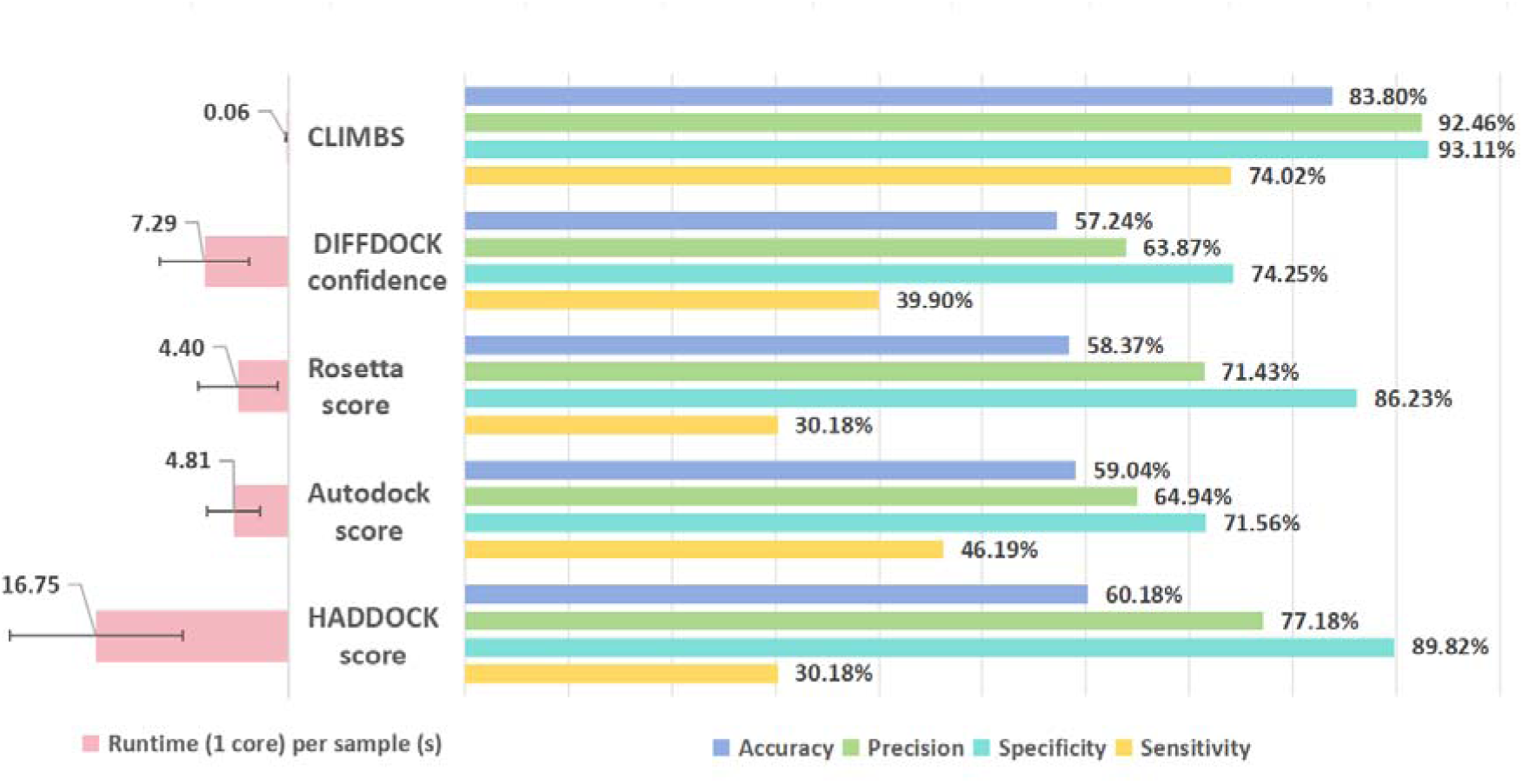
Comparison between CLIMBS and other methods by different indicator: runtime per sample, accuracy: (TP+TN)/(TP+FP+TN+FN), precision: TP/(TP+FP), specificity: TN/(TN+FP) and sensitivity: TP/(TP+FN). TP True Positive, TN True Negative, FP False Positive, FN False negative. Thresholds of predicting as bound for each metric are in Supplementary Data 8.

CLIMBS spends only 0.06s to predict one sample in one CPU. It is mainly because the classifier only considers the residues around the protein-sugar interface. CLIMBS has the best accuracy of 83.80% and precision of 92.46%, which is more than 15% better than other methods compared. It also has a remarkable sensitivity of 74.02%, while other methods are under 40%. All five methods have higher specificity than sensitivity, indicating that methods perform better on negative samples than positive samples.

CLIMBS has the smallest difference of 19.02% between specificity and sensitivity.

We have then investigated the methods’ performance using the Wasserstein distance to describe the difference between distributions of positive and negative samples (Figure 3). CLIMBS’ predictions are separated, with 0.670 Wasserstein distance. For the other four methods, the positive and negative distributions are overlapped in most cases, with 0.049 being the highest Wasserstein distance. Thus, CLIMBS can distinguish well between bound and non-bound samples within a diverse sugar binder dataset.

**Figure 3.**
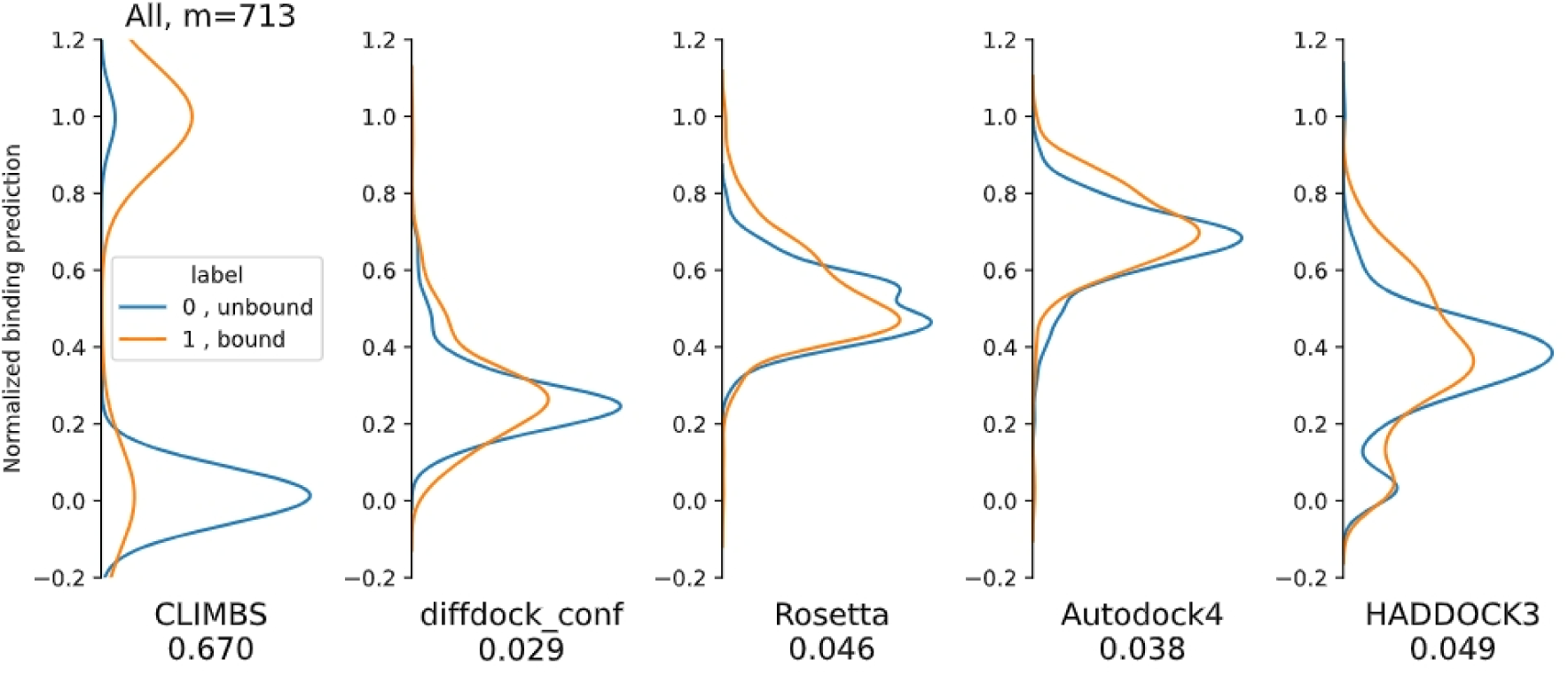
Comparison between CLIMBS and other methods by Wasserstein distance across multiple carbohydrates. Scores from each method were normalised between 0 and 1, corresponding to the lowest and highest scoring samples. Distributions of normalised scores are plotted as kernel density estimates. Positive samples are shown in orange and negative samples in blue. The number under each subplot is the Wasserstein distance between the normalised score distribution of positive/negative samples for each method. m: tested sample amount. For the dataset see db_eval Supplementary Data 3j.

### Sugar-type specificity

Due to the structural similarity of mono-saccharides, interaction specificity is an important performance indicator in modelling sugar-protein interactions. Therefore, we have evaluated CLIMBS specificity performance on the carbohydrate-binding domain of *Marinomonas primoryensis* PA14 (MpPA14), which binds to a series of mono-saccharides except N-acetyl-galactosamine (GalNAc). The apparent dissociation constants (K_d_app) of MpPA14-sugar complexes were measured by isothermal titration calorimetry (ITC) in previous work. (*32*)

Six solved complexes (*32*) and one putative structure with GalNAc (generated by aligning and replacing GlcNAc with GalNAc in a solved MpPA14-GlcNAc complex) were selected as initial structures after Rosetta relaxation. Each initial structure underwent a Rosetta local docking protocol (*33*) that generated 200 docked structures. CLIMBS was used to evaluate the binding of the initial and the docked structures.

Results in Table 2 show that the classifier, similarly to Rosetta Energy Function, can correctly identify that GalNAc is not binding while the other sugars are bound. However, for the best binding complex with fucose, Rosetta Energy Function over-estimates the binding free energy.

**Table 2.** Binding between MpPA14 and different sugars. Previous work (32) measured the apparent dissociation constants (K_d_app) by isothermal titration calorimetry, and the binding free energy (ΔΔG) was calculated based on K_d_app with a temperature of 298 K. Rosetta Energy Function and CLIMBS were used to estimate the binding of relaxed solved structures and the structure after running a Rosetta local docking. The local docking test values are the average and standard deviation of the results of 200 docking runs. For CLIMBS the values represent the fraction of times that docking poses have been scored as positive. PDB ID of initial structure: 6X7J, 6X7Y, 6X8A, 6X7X, 6X8D, 6XAC.

| Sugar | Experiment verification |  | Crystallized structure |  | Rosetta local docking test |  |
| --- | --- | --- | --- | --- | --- | --- |
| | Kdapp(mM) | $\Delta\Delta G_{298K}$ (kcal/mol) | CLIMBS | Rosetta $\Delta\Delta G$ (kcal/mol) | CLIMBS | Rosetta $\Delta\Delta G$ (kcal/mol) |
| L-Fucose | 0.65 | -4.33 | 1 | -7.08 | 0.90 | -9.60±0.76 |
| N-acetyl-glucosamine | 1.07 | -4.04 | 1 | -4.53 | 1.00 | -3.34±1.27 |
| Glucose | 1.4 | -3.88 | 1 | -4.95 | 0.90 | -3.07±0.91 |
| Mannose | 1.7 | -3.76 | 1 | -3.75 | 1.00 | -2.37±0.77 |
| L-Arabinose | 2.2 | -3.61 | 1 | -4.56 | 0.98 | -0.03±0.57 |
| Galactose | 4.1 | -3.24 | 1 | -2.88 | 0.98 | -1.89±0.89 |
| N-acetyl-galactosamine | -- | -- | -- | -- | 0.30 | 4.81±1.63 |

### Medium-trained model to predict binding of unseen sugars

Several types of sugars have no solved structure in complex with binding protein and CLIMBS is likely to face samples that include sugar types not used for training. To test the performance on unseen carbohydrates, a model (*db_p3,* see Supplementary Data 3d) trained on samples including various sugars except for GlcNAc-GlcA (NAG-BDP), arabinose (ARA), 5-O-phosphono-ribose (RP5) and 2-N-sulfo-6-O-sulfo-D-glucosamine (ASG) was used to test four unseen sugars with different training epochs (Figure 4). As the training epoch increases, the validation loss of trained sugars samples converges to 0.08 at 400 epochs. The loss of arabinose samples performs like trained samples losses, decreasing along with the training. However, the losses of the other untrained sugars samples stop converging between 100 to 200 epochs. And the loss of the untrained di-saccharide even begins increasing and reaches 0.29 at 450 epochs while 0.13 at 275 epochs. The diverging losses of training and untrained sets indicate that overfitting of the model happens as the training epoch increases, indicating that a medium-trained model between 200 to 300 epochs) is therefore an appropriate choice for predictions of untrained carbohydrates.

**Figure 4.**
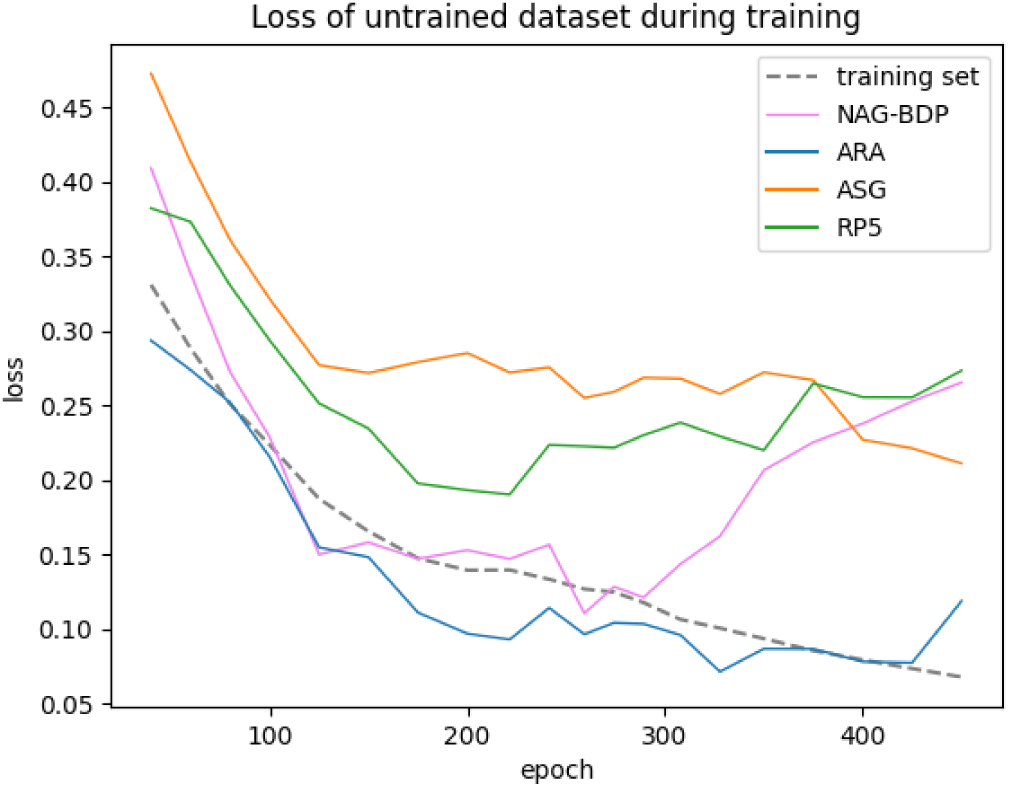
Loss curves of trained and untrained samples during CLIMBS training (see Supplementary Data 3d). The training set included 14 types of sugar binders (see table 1). Four types of untrained samples are GlcNAc-GlcA (NAG-BDP), arabinose (ARA), 5-O-phosphono-ribose (RP5) and 2-N-sulfo-6-O-sulfo-D-glucosamine (ASG) binder.

### Retraining the model for new sugars

CLIMBS can be retrained for new sugar types with limited number of samples, which we assessed with a series of retraining tests.

An initial database with samples containing proteins in complexes with highly represented carbohydrates in the PDB (Glc, GlcA, GlcNAc, galactose, fucose, mannose and xylose in *db_r0*, see Supplementary Data 3e) was used to train a starting model.

These seven sugars contain common shared fragments (e.g. central ring and hydroxyl groups). Our hypothesis was that, due to shared overall structures, accuracy for new carbohydrate could be rapidly increased with only a few samples introducing specific features. Pairs of additional carbohydrates with similar features and absent from the initial dataset were selected: one was added to the initial database for training and validation, and the second used for monitoring accuracy improvement without training.

Two pyranoses, sialic acid and arabinose, were selected as first candidates with starting accuracy at 67.57% and 78.26%, respectively (Figure 5A). The lower value for sialic acid might be due to a larger structural complexity (nine-carbon backbone) and the absence of an atom type (CH0 atom) in the database, while all atom types for arabinose are already present in the database. Sialic acid samples were progressively added to the training set, while arabinose was kept untrained (*db_r1,* see Supplementary Data 3f). For sialic acid, adding 12 samples is sufficient to reach an approximate 80% accuracy, without decreasing the control accuracy (arabinose). The control accuracy dropped when more than 12 samples were added, indicating overfitting.

**Figure 5.**
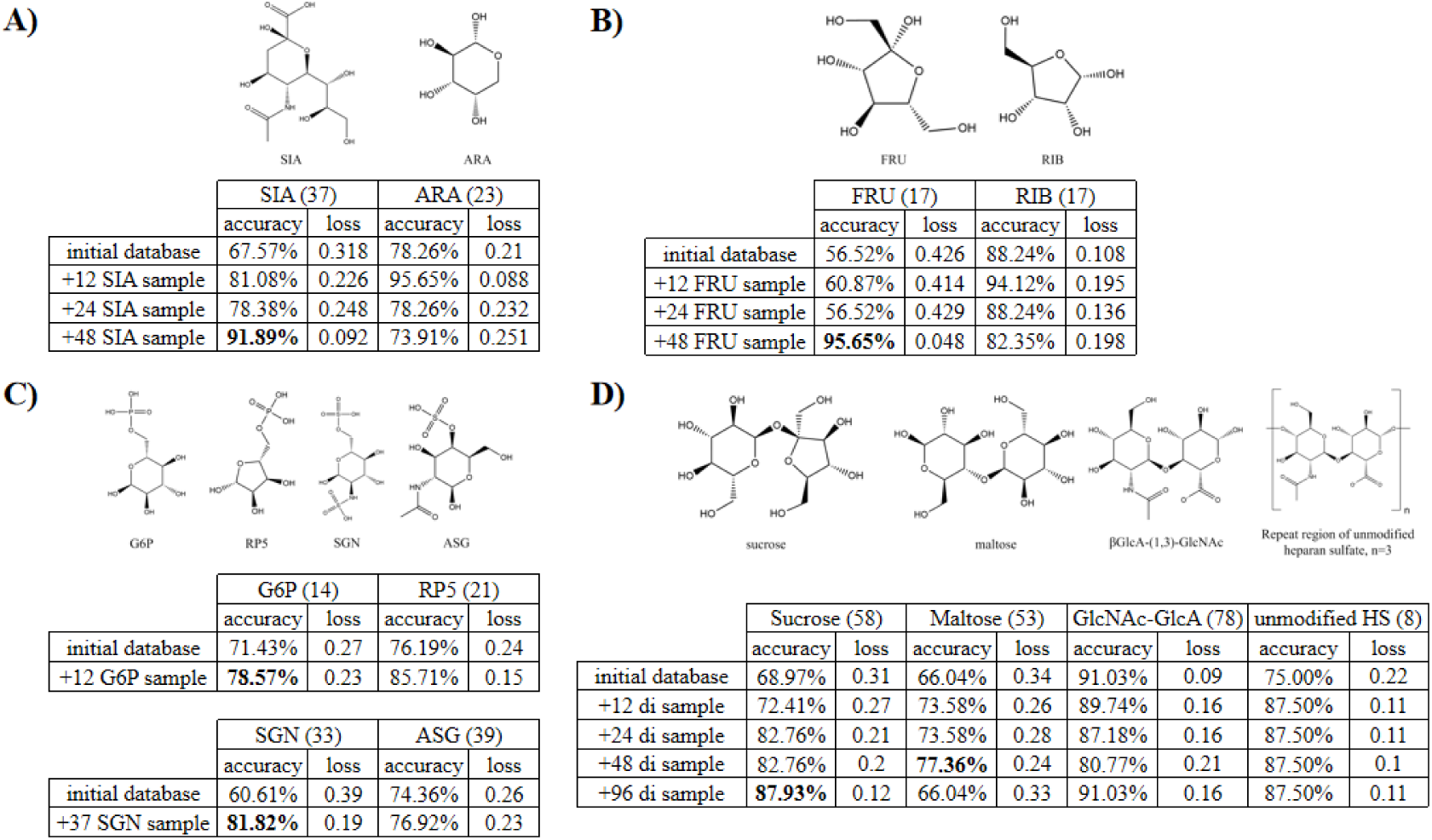
Retraining CLIMBS with new sugars. A. Accuracy and training loss of arabinose (ARA, 23 samples in test set) and sialic acid samples (SIA, 37 samples in test set) from a CLIMBS model trained on initial database plus different number of sialic acid samples (see Supplementary Data 3f). B. Accuracy and training loss of ribose (RIB, 17 samples in test set) and fructose (FRU, 17 samples in test set) samples from a CLIMBS model trained on initial database plus different number of fructose samples (see Supplementary Data 3g). C. (Above) Accuracy and training loss of 6-O-phosphono-glucose (G6P, 14 samples in test set), 5-O-phosphono-ribose (RP5, 21 samples in test set) from a CLIMBS model trained on initial database plus different number of 6-O-phosphono-glucose samples. (Bottom) Accuracy and training loss of N,O6-disulfo-glucosamine (SGN, 33 samples in test set), N-acetyl-4-O-sulfo-galactosamine (ASG, 39 samples in test set) and trained. D. Accuracy and training loss of maltose, sucrose, GlcNAc-GlcA and repeat regions of unmodified heparan sulphate samples from a CLIMBS model trained on initial database plus different number of maltose and sucrose samples (See Supplementary Data 3i). The ratio of maltose and sucrose is 1:1. HS: heparan sulphate.

**Figure 6.**
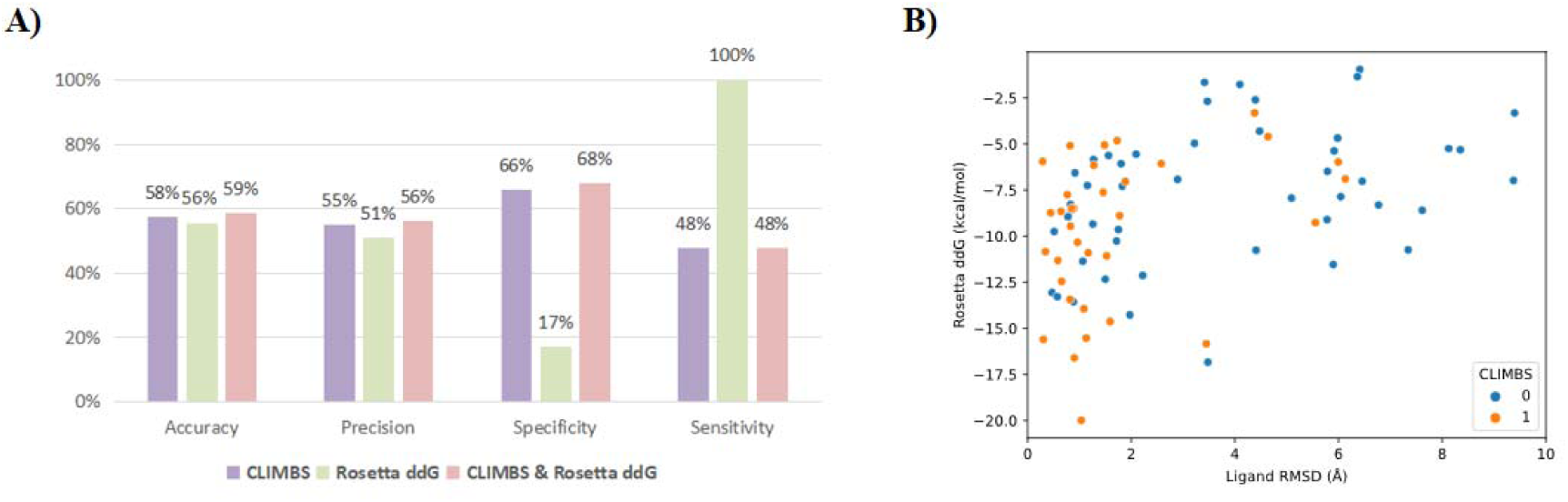
Scoring methods on carbohydrate-protein docking problem. 99 samples were included. Structures with ligand RMSD <2 Å after superimposing the protein to the referral structure are regard as bound. A) Performance of each method. B) detail scatter plot for each carbohydrate-protein complex sample. Thresholds to predict as positive in for each metric: CLIMBS=1, Rosetta score function ddG <-4.5. For performance with different threshold Supplementary Data 9.The positive predictions of combined metrics are the AND subset of each positive predictions. accuracy: (TP+TN)/(TP+FP+TN+FN), precision: TP/(TP+FP), specificity: TN/(TN+FP), sensitivity: TP/(TP+FN). TP: True Positive, TN: True Negative, FP: False Positive, FN: False Negative.

As the second case, we introduced fructose and ribose (Figure 5B), which both contain a furanose ring not seen in the initial database (all trained carbohydrate types have a pyranose ring). The initial binding accuracy was high for ribose (88.24%) but not for fructose samples (56.52 %). The lower value for fructose might be due to the absence of an atom type (CH0 atom) in the database. Then, fructose samples were added to the training set (*db_r2,* see Supplementary Data 3g). The accuracy of fructose samples increases significantly at 48 samples to 95.75%, however showing a small decrease in ribose accuracy.

We also investigated how many samples were needed to retrain CLIMBS for new monosaccharides that include chemical modifications (Figure 5C). Phosphorylation and sulphonation of carbohydrates are two frequently observed functional modifications that were not present in the initial database. N,O6-disulfo-glucosamine (SGN) and 6-O-phosphono-glucose (G6P) were selected for model training and testing, while N-acetyl-4-O-sulfo-galactosamine (ASG) and 5-O-phosphono-ribose (RP5) only for testing (*db_r3,* see Supplementary Data 3h). Surprisingly, the initial model wasn’t trained on phosphate and sulphate has accuracies between 60% and 76% to predict binding of modified sugars, likely driven by the rest of the carbohydrate structural features. Retraining with less than 40 samples with phosphorylated/sulphonated sugar improved binding accuracy to 76%-85%, both for trained and untrained carbohydrates.

At last, we investigated retraining for di-saccharides constructed by seen monosaccharides (Figure 5D). Maltose (glucose-α-1,4-glucose) and sucrose (glucose-α-1,2-fructose) were selected as candidate di-saccharide for training and testing (*db_r4,* see Supplementary Data 3i). Additionally, a heparan sulphate fragment, GlcNAc-GlcA (β-1,4 and β-1,3 linkage) disaccharide unit was added for testing. The initial model, trained on only monosaccharides samples, has considerable performance on the disaccharide structures, despite no glycosidic bond being included in the training set. The accuracy of sucrose and maltose increases from near 70% to 88.93% with 96 extra samples for sucrose and to 77.36% with 48 extra samples for maltose. The accuracy of GlcNAc-GlcA and repeat regions of unmodified heparan sulphate are hardly affected and fluctuates at high value around 90%. It shows that the model trained by monosaccharides is able to assess disaccharide and even polysaccharide accurately. More generally, introducing 48 disaccharide samples into the training set can increase accuracy before overfitting.

### Application to protein docking

CLIMBS can practically work as an additional scoring method on carbohydrate-protein complex modeling problems. We first applied it to protein-carbohydrate global docking, to predict correct placement of the ligand for complexes with solved crystal structures in dataset *db_dock* (Supplementary Data 3k), which included 197 samples with resolution<2Å.

Docked conformations were generated with Chai-1(*34*). Only for 99 samples, the top scoring model passed the structure quality filters (see Supplementary Data 9) and these complexes were used for docking assessment.

We defined as true positive docks the poses with ligand RMSD <2 Å after superposition of the protein model to the solved structure. CLIMBS, Rosetta score and a combined score CLIMBS + Rosetta were used to evaluate the binding state of the predicted structures. Models were considered positive (bound) for Rosetta score <-4.5 kcal/mol, capturing the majority of interactions observed in crystal structures for protein-carbohydrate complexes (see Supplementary Data 4). The CLIMBS + Rosetta method scores a structure as bound when both CLIMBS and Rosetta score it as bound, and evaluates it as non-bound when either CLIMBS or Rosetta scores it as non-bound.

CLIMBS shows accuracy of 58%, slightly higher than Rosetta score. The CLIMBS + Rosetta combined score performs similarly to CLIMBS alone, indicating that they are discriminating complexes in similar way. Within this test set, Rosetta’s sensitivity rises to 100% because no false negative samples are present with ddG >-4.5 kcal/mol cut off, equivalent to Kd=1 mM. Stricter energy cutoffs, such as-8 kcal/mol and-13 kcal/mol improve accuracy for our test set (see Supplementary Data 9) but might be too strict for likely carbohydrate binders that operate in the low mM to high μM range.

### Application to protein design

We then assessed CLIMBS and Rosetta on evaluation of a protein design task. As test set, we employed engineered carbohydrate-binding proteins developed through directed evolution from a non-binding scaffold (*35*). Binding affinities and specificities have been experimentally determined for 4 variants and 13 carbohydrates, but no crystal structure is available. Complexes matching the experimental specificities were labelled as positives (bound), while others were labelled as negatives (unbound) (see Supplementary Data 10). All 52 complexes were predicted with Chai-1 (*34*) and the top prediction was considered. The top prediction of only 37 generated structures passed the structure-quality filter (see Supplementary Data 9) and were evaluated with CLIMBS, Rosetta and the CLIMBS + Rosetta score (Figure 7). CLIMBS shows 70% overall accuracy, while Rosetta reaches only 24%. CLIMBS + Rosetta shows the same accuracy as CLIMBS. Stricter cutoffs on the Rosetta score did improve the performance, resulting in 65% accuracy for Rosetta score, and 81% for CLIMBS + Rosetta score (see Supplementary Figure 5). Overall, CLIMBS shows a reliable performance on the design problem, and the combination with Rosetta can improve the overall accuracy.

**Figure 7.**
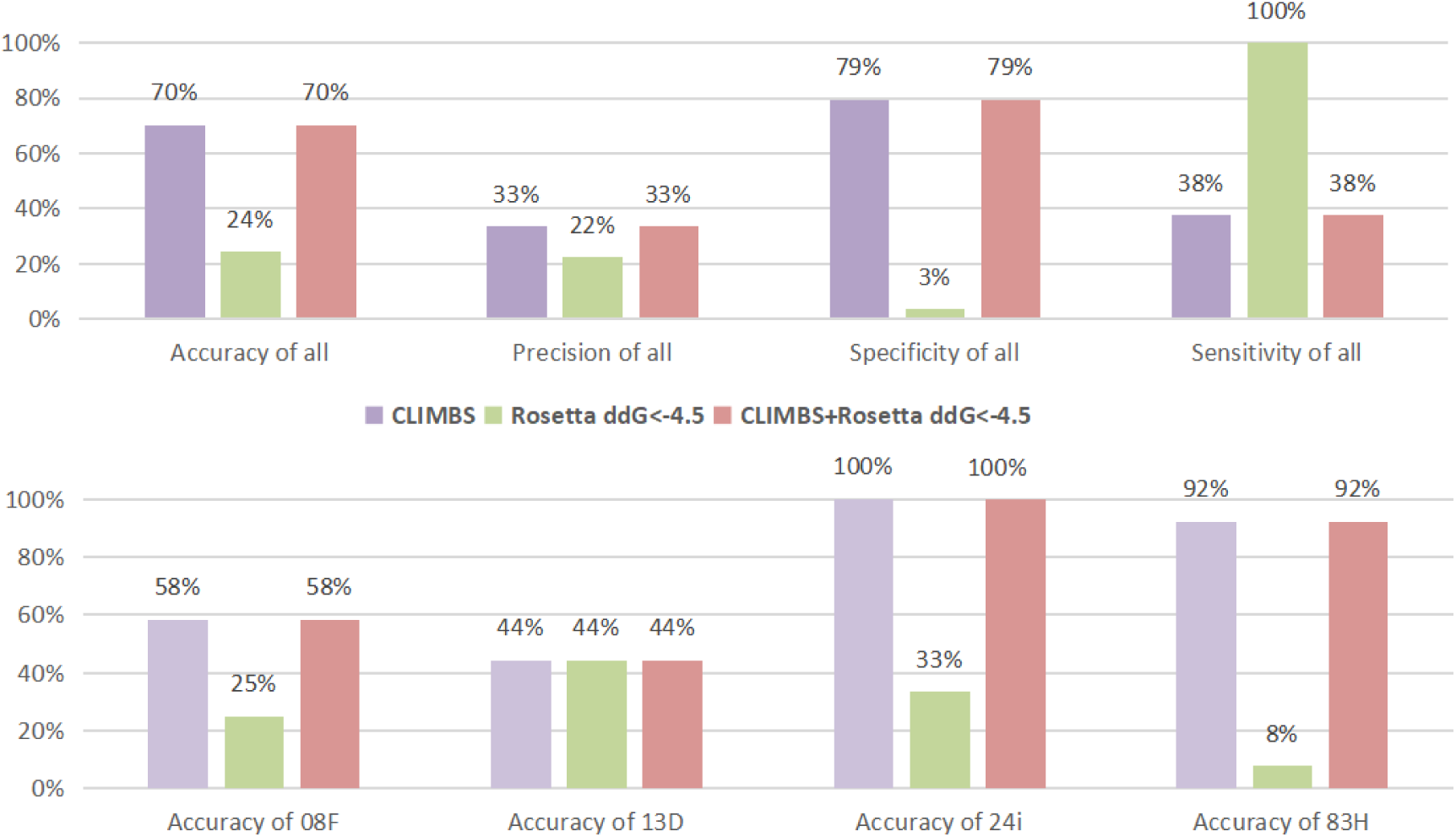
Performance of different scoring methods on design carbohydrate-binding protein problem. In total 37 samples were included: 12 samples of 08F, 9 samples of 13D, 3 samples of 24i, 13 samples of 83H. 08F, 13D and 24i are binders with different specificities, 83H does not bind to any of the 13 targets analyzed. Four protein variants bind to different carbohydrate in 13 candidates. Thresholds to predict as positive in for each metric: CLIMBS=1, Rosetta score function ddG <-4.5. For performance with different threshold Supplementary Data 9. The positive predictions of combined metrics are the AND subset of each positive predictions. accuracy: (TP+TN)/(TP+FP+TN+FN), precision: TP/(TP+FP), specificity: TN/(TN+FP), sensitivity: TP/(TP+FN). TP: True Positive, TN: True Negative, FP: False Positive, FN: False Negative

## IV. Discussion

In this study we have developed CLIMBS, a machine learning classifier that assess the structure of binding pockets in carbohydrate-protein complexes models.

A sugar-protein complex library was built to include 21 different carbohydrates for model training and testing. Solved sugar-protein complexes and failing designed carbohydrate binders were collected and labelled as positive and negative samples, while other methods tend to use solved binder structures with scalar labels such as experimental binding affinity. A binary label was chosen because there is not a scalar experimental result for non-binding and, within a protein design pipeline framework, a primary focus would be identifying non-binding complexes that should not be taken into the next steps.

On one hand, this means that CLIMBS cannot describe the strength or extent of sugar-protein interactions, although, as in the case of validation of multiple local docking poses, a crude but effective scalar value can be established by averaging the scores of the entire population. On the other hand, CLIMBS focuses on effectively discriminating good-binding and poor/non-binding complexes, using a method independent from and unrelated to currently used score functions or confidence metrics.

The use of negative data originated from a computational protein design pipeline, is a distinct feature of our approach to classifier training, showing the power of synthetic data augmentation to overcome the limitations of current scoring functions, which were used to design these structures in first place.

The choice of suboptimal carbohydrate binder designs as synthetic negative data, was a key element in the setup of the training dataset and in the performance of CLIMBS. It allowed us to build a balanced dataset and to focus training on atomic details specific to carbohydrate protein interactions. CH-π interactions, which are often fundamental for sugar binding but not factored in by most energy-based methods, are included as a key type of interaction in this model.

A fragment-level representation keeps the necessary information among the functional groups of atoms, facilitating learning of amino acid features that are present in multiple molecules (e.g. the carboxyl group in aspartic acid and glutamic acid).

CLIMBS generalizable approach to protein carbohydrate interaction allows to address sugars not included in the training set. Lower epoch models can be used to avoid potential overfitting, or a few tens of new samples can be added to the dataset to retrain. These characteristics make CLIMBS architecture particularly suitable for carbohydrate-protein investigation, since databases are relatively small and different carbohydrates have similar structures, enhancing the recognition of these common features.

This versatility is extended also to carbohydrate binding sites that contain metal ions, as for the MpPA14 used for assessing carbohydrate specificity, which contains a calcium ion. 141 carbohydrate-protein complexes in the library contain at least one metal atom in the binding site, which is considered implicitly in CLIMBS; although the metal ions were removed during the data preprocessing, the residues interacting to the metal ions were kept and provided with this information during training. As a result, CLIMBS can work with carbohydrate-protein complexes that include metal ions at the interface.

Compared to other score functions and machine learning methods, CLIMBS has shown outstanding overall accuracy and precision, fast runtime, and better performance on recognition of bound samples. As an entirely geometry-based method, it can distinguish different sugars with a highly similar structures that energy-based methods cannot. It is therefore a new scoring tool, orthogonal to other energy and machine learning based methods, that, when combined with docking and design protocols and pipelines, allows to improve selection of protein-carbohydrate models that retain native-like features. This is particularly relevant in comparison with Rosetta, as training was performed on Rosetta-relaxed structures. The sub-second runtime makes it feasible to combine with current energy-based methods or machine learning methods, filtering false positive/negative results and increasing the overall pipeline accuracy.

On practical applications such as scoring of docking and design, CLIMBS’ accuracy decreases, as expected, since the tasks differ slightly from the training sets and are more challenging. CLIMBS was trained to distinguish native bound structures and artificially-generated non-bound structures, although those are often energetically similar. However, in realistic docking and design cases, positive and negative samples are usually not known, and they are produced using the same method (generative model followed by Rosetta relax in our case). The quality of samples passed to CLIMBS is also critical: only about half of the generated models were acceptable in terms of quality metrics (protein and ligand pLDDT) and this can bias the range of complexes analysed, with a chance of potentially loosing weaker binders.

Training CLIMBS on a restricted dataset containing only high-resolution structures (<2) (dataset *db_w2*, Supplementary Data 3l) resulted in a model able to provide higher discrimination (64% accuracy up from 58%) in docking assays where the test set was also composed only of high-resolution structures of complexes, likely to have high affinity.

However, in practical cases where binding affinity is likely in the millimolar or high micromolar range, like in the design cases, the general model still performs best, and the performance can be further boosted by combining it with Rosetta scoring.

CLIMBS is characterized by architectures and embeddings that leverage shared structural features of carbohydrates, and by training with negative synthetic data that have driven its discrimination power. This approach can be potentially generalized to other supervised learning methods, making use of this large untapped resource of data.

## V. Methods

### Model architecture

CLIMBS takes a PDB structure file containing a carbohydrate chain and one or more protein chains as input, and it outputs a value of 1 (binding) or 0 (not binding). A protein-sugar interacting graph is generated from data pre-processing. Then, the interacting graph is passed to the embedding module including spherical message passing layers (*36*), principal neighborhood aggregation layers (*37*), and a super-node-based hierarchical graph pooling layer (*38*). Followed by fully connected layers and a sigmoid function, a binary result is returned.

Data pre-processing elicits the essential details of protein-sugar interactions in the pdb structure file (see Supplementary Data 7). Firstly, it selects the amino acid and carbohydrate residues on the interface. Only the amino acid/saccharides that have polar or CH-π interaction to the saccharide/amino acids are kept (see Supplementary Data 1). Hydrogen atoms are discarded. Secondly, it elicits the minimum binding site from the interface residues: for amino acids, either the terminal functional groups of sidechains or the mainchain backbones are kept; for saccharides, all heavy atoms are kept. Finally, the information of the minimum binding site is generated, including atom coordinates, an atom-level interacting graph *G={V,E}* and an assignment matrix *S*. Node matrix *V* includes the one-hot encoded atom types. Edge matrix *E* describes three types of interaction between nodes: chemical bond, polar interaction and CH-π interaction.

Assignment matrix *S* assigns atoms into corresponding fragments. Fragments are groups of atoms that commonly exist in different molecules. For example, glucose is constructed by a 6-member ring fragment, 4 oxygen fragments and a carbon-oxygen (single bond) fragment. The fragment-level pooling layer of CLIMBS can capture the features of fragments shared among different types of sugars.

Given the pre-processed data, the classifier model returns a binary value indicating bound or not (see figure 5). In the beginning, the interaction graph *G={V,E}* as well as the atom coordinates are passed to spherical message passing (SMP) layers (*36*) to generate an updated graph *G’={h,E}.* The first SMP layer converts the atom coordinates into three geometrical representations (distances, angles, and torsions) and incorporates them into the interaction graph *G*. Then, the rest of SMP layers update edges and aggregates them into node feature matrix *h*. Then, principal neighbourhood aggregation (PNA) (*37*) layers update *G’={h,E}* into *G’’={h’,E}* by a multiple aggregation function and degree-scalers.

After that, a pooling layer coarsens the atom-level graph *G’’={h’,E}* into a fragment-level graph *G*={h*,E*}* with the assignment matrix *S*. Then, the fragment-level node feature *h\** is passed to a sum aggregator and an activation function to output a global feature *u*. The global feature *u* is then passed to fully connected layers and a sigmoid function to output a normalized prediction *p* [0,1]. Finally, 0 or 1 is outputted by comparing the *p* to a threshold value (default 0.5, for detail see Supplementary Data 5) to indicate as unbound or bound.

### Datasets

Positive samples were derived from the Protein Data Bank. Structures were validated (see Supplementary Data 4) to ensure high quality structures are used. To increase reliability and accuracy, redundance cleaning was done for the positive database according to protein-sugar complex RMSD and interface RMSD. All positive samples were relaxed by Rosetta (*39*) to alleviate potential clashes.

Negative samples were obtained from computational design. Experimental success rate of designed binders for small molecule is around 0.07% according to the latest RFAA design method (*27*), though with high scoring for energy-and structure-based metrics, e.g. AF2 pLDDT > 80, designed RMSD <2 Å low Rosetta ddG (<-30 in the case of digoxigenin binder) and at least one hydrogen bond to ligand. Therefore, poorly scoring designs can be reasonably assumed as failures. Here, A Rosetta design protocol (unpublished) was used to generate carbohydrate-protein complexes. The top 10% worst (ranked by binding free energy of Rosetta Energy Function) complexes were selected as negative samples. In general, those sugar binders underperform because of either lack of interactions (e.g. cavities, shallow clefts), or inaccurate interactions (e.g. clashes and distorted hydrogen bonds).

### Model optimization

The training of CLIMBS was supervised by the binary 0/1 label of negative and positive samples. Mean absolute errors (MAE, also known as L1 loss) between the label and the prediction *p* were used for parameter optimization. The sugar-protein complex structure library comprises a ratio of positive and negative samples which is approximately 1:1. 15 types of mono-saccharides, 3 types of di-saccharides and a linear poly-saccharide with the corresponding binders, in total 13507 samples are in the library (see Supplementary Data 3a). The library was split into training, validation and test sets with the ratio of 2:2:1, while maintaining the 1:1 positive and negative ratio (for training detail see Supplementary Data 2).

### Pooling layer selection

We have explored three representations of minimum binding site as pooling layers according to the scale of molecular structure detail: atom-level, fragment-level and molecule-level representations. Classifier models with those three different pooling layers before the sum aggregator were trained and tested (results in Supplementary table 3), with the fragment-level pooling layer performing best for both mono-saccharide prediction and di-saccharide prediction.

### Robustness check

A robustness check was performed to unravel the factors contributing to CLIMBS’ performance. Model robustness is the ability of the model to maintain performance when faces adversarial data compared to the training data. Adversarial samples were generated by slightly modifying the original samples such as with masked edges and extra nodes or edges in the interacting map G. Results in Supplementary table 4, reveal that CH-π interacting residues increase the probability of prediction as positive (bound), but it is not the only factor, consistent with experimental results from Hudson (*14*).

## VII. Acknowledgements

Yijie Luo is under support of China Scholarship Council. Fabio Parmeggiani is the recipient of an EPSRC early career fellowship (EP/S017542/2). The authors thanks Dek Woolfson and the Woolfson group for the support and the stimulating discussions.

## VIII. Author Contributions

The research project was conceived by Yijie Luo (YL) and Fabio Parmeggiani (FP). YL developed the code and collected the data. FP supervised the project. YL and FP analysed the data and wrote the manuscripts.

## IX. Competing interests

The authors declare no competing interests.

## X. Data and materials availability

The code and CLIMBS model are available as part of the Rosetta ML package (https://github.com/Parmeggiani-Lab/sugar_binding_predictor2).The dataset of positive and negative samples is available in Zenodo (https://zenodo.org/records/15169999). The protocol used to design negative sample is documented in https://github.com/Jacky233emm/new_residue_design.

## Notes

### Competing Interest Statement

The authors have declared no competing interest.

### Summary of Updates

revision of text, additional results on docking and design tests

