## supplementary information for "CLIMBS: assessing carbohydrate-protein interactions through a graph neural network classifier using synthetic negative data"

**Affiliations:**

### Supplementary Data 1. Definition of polar interaction and CH-π interaction

Polar interactions between molecules were described as the non-carbon heavy atom pair within 3.5 Å. CH-π interactions were defined according to previous research (*1*). In short, a CH-π interaction exists between the CH group of the sugar ring and aromatic amino acids, limited by the distance (<4.5Å) of C to the center of the aromatic ring, the angle between the CH vector and the aromatic ring normal (<40°), and the distance between the C projection on the aromatic ring and center of the aromatic ring (<2Å).

### Supplementary Data 2. Model optimization

The dataset for model optimization was split into training, validation and test sets with the ratio 2:2:1. The model in which validation loss was lower than training loss was regarded as having completed training. The maximum training epoch was 500. The performance (e.g. accuracy and specificity) of trained models was tested by the test set.

Each negative binder sample generated by Rosetta(*2*) includes a CH-π interacting residue to the binding sugar. However, only around 30% of the native binders from RCSB(*3*) (positive sample) had CH-π interaction to sugars. Pre-testing had shown that using original negative samples for training would lead to severe bias of CH-π interaction. Thus, only 30% of the negative samples would keep the CH-π interacting residue during data pre-processing.

During model training, merged samples were used to reduce the bias between binding complexes including mono-saccharide and more than one carbohydrate unit. Merged samples were generated by combining two different structures files with mono-saccharide into one structure file. Only all-positive or all-negative merged samples were used.

### Supplementary Data 3. Dataset detail for used models

This section states the components of the datasets for the models used. Complexes with 23 different ligands were collected in the library(*4*) (supplementary table 1).

Supplementary table 1 Ligand type involved in the library of carbohydrate-protein complex.

| **Carbohydrate type in complex** | **Ligand ID** |
| --- | --- |
| N-acetyl-α-D-galactosamine | A2G |
| α-L-arabinose | ARA |
| N-acetyl-4-O-sulfo-β-D-galactosamine | ASG |
| β-D-glucuronic acid | BDP |
| 6-O-phosphono-β-D-glucose | BG6 |
| β-D-glucose | BGC |
| GlcNAc-(1,4)-βGlcA & βGlcA-(1,3)-GlcNAc | di12 |
| maltose | di3 |
| sucrose | di4 |
| β-D-fructose | FRU |
| α-L-fucose | FUC |
| 6-O-phosphono-α-D-glucose | G6P |
| β-D-galactose | GAL |
| α-D-glucose | GLC |
| α-D-mannose | MAN |
| N-acetyl-β-D-glucosamine | NAG |
| N-acetyl-β-D-galactosamine | NGA |
| Repeating region of heparan sulphate  (constructed by 3 GlcA and 3 GlcNAc) | poly1 |
| α-D-ribose | RIB |
| 5-O-phosphono-β-D-ribose | RP5 |
| N,O6-disulfo-glucosamine | SGN |
| N-acetyl-α-neuraminic acid | SIA |
| α-D-xylose | XYS |

As mentioned, there are training, validation and test sets in the model optimization dataset. Some models used extra datasets to evaluate the properties of the model. Samples in one type of ligand were used for training/validation/test sets with a ratio of 2:2:1 in model optimization, for validation/test set a ratio 2:1 in extra evaluation, or for 100% test set in testing only.

#### a. Dataset for final model (*db_w1*):

Complexes with ligand A2G, BDP, BG6, BGC, di3, di4, FRU, FUC, G6P, GAL, GLC, MAN, NAG, NGA, SGN, SIA and XYS were used for model optimization. Complexes with ligand RIB, ARA, ASG, di12, poly1 were for testing only.

#### b. Dataset for pooling layer optimization (*db_p1*):

Complexes with ligand NAG, BDP and di12 were used for model optimization.

#### c. Dataset for robustness check (*db_p2*):

Complexes with ligand NAG and BDP were used for model optimization. Complexes with ligand GLC were for testing only.

#### d. Dataset for predicting unseen carbohydrates (*db_p3*):

Complexes with ligand GLC, NAG, BDP, BGC, GAL, FUC, MAN, SIA, XYS, FRU were used for model optimization. Complexes with ligand di12, ARA, ASG, RP5 were for testing only.

#### e. Dataset for new sugars retraining (initial, *db_r0*):

Complexes with ligand GLC, NAG, BDP, BGC, GAL, FUC, MAN, XYS were used for model optimization.

#### f. Dataset for new sugars retraining (unseen mono-saccharide, *db_r1*):

Complexes with ligand GLC, NAG, BDP, BGC, GAL, FUC, MAN, XYS were used for model optimization. Complexes with ligand ARA, SIA were used for extra evaluation. Different number of SIA (0,12,24,48 samples) samples are added in training set.

#### g. Dataset for new sugars retraining (unseen furanose ring, *db_r2*):

Complexes with ligand GLC, NAG, BDP, BGC, GAL, FUC, MAN, XYS were used for model optimization. Complexes with ligand RIB, FRU were used for extra evaluation. Different numbers of FRU (0,12,24,48 samples) samples are added to the training set.

#### h. Dataset for new sugars retraining (unseen fragment, *db_r3*):

Complexes with ligand GLC, NAG, BDP, BGC, GAL, FUC, MAN, XYS were used for model optimization. Complexes with ligand RP5, G6P, ASG, SGN were used for extra evaluation. Different numbers of G6P (0,32 samples) as well as SGN (0,37 samples) samples are added to the training set.

#### i. Dataset for new sugars retraining (unseen di-saccharide, *db_r4*):

Complexes with ligand GLC, NAG, BDP, BGC, GAL, FUC, MAN, XYS were used for model optimization. Complexes with ligand di12, di3, di4 were used for extra evaluation. Different number of di3 (0,6,12,24,48,53 samples) as well as di4 (0,6,12,24,48,60 samples) samples are added in training set.

#### j. Dataset for methods’ comparison (*db_eval*):

Here is the final model (Supplementary Data 3a). Dataset for model performance evaluating samples in testing set of NAG, GLC, BGC, GAL, FRU, di3, di4 after splitting.

#### k. Dataset for docking application evaluation (*db_dock*):

It includes 197 samples in testing set with PDB structure resolution<2 Å, containing ligand XYS, SIA, NGA, NAG, MAN, GLC, GAL, G6P, FUC, FRU, BGC, BG6, BDP, ASG, ARA, A2G.

#### l. Dataset for re-training model to improve performance on docking application (*db_w2*):

High PDB structure resolution (<2 Å) complexes with ligand A2G, BDP, BG6, BGC, di3, di4, FRU, FUC, G6P, GAL, GLC, MAN, NAG, NGA, SGN, SIA and XYS were used for model optimization. Complexes with ligand RIB, ARA, ASG, di12, poly1 were for testing only.

### Supplementary Data 4. Processing positive samples and generating negative samples

Positive samples were experimentally solved structures of native sugar binding proteins from the Protein Data Bank. Structures with good quality were kept. Structure quality was assessed by PDB resolution (<3Å), atom occupancy (>0.9) and B-factor (<80) (*5*). Ligand structure quality was further assessed by RSCC (>0.8), RSR (<0.3), bond RMSZ (<2) and angle RMSZ (<2) (*6*). Each target sugar and nearby (within 4.5Å for aromatic residue and 3.5 Å for other) protein chain(s) were saved as a sugar-protein complex. Complexes with same ligand, sequence identity >25% and binding site (sugar around 4.5Å) RMSD <1Å were regarded as redundant, only one of them would be kept. Then, the complexes were relaxed with coordination-constrained by Rosetta. To strip out the samples with low binding affinity, filters including interface residue number >4 and Rosetta ddG <-4.5 kcal/mol (approximately dissociation constant 0.5 mM). The ΔΔG cutoff is closed to the binding free energy value in native carbohydrate binding protein (*7*), and also near to the peak of Rosetta ddG distribution of positive sample before filtering.

Supplementary figure 1 Rosetta ddG distribution of positive samples after Rosetta relaxation and before filtering. 90% of sample has ddG from -0.69 to -19.69 (allowance range).

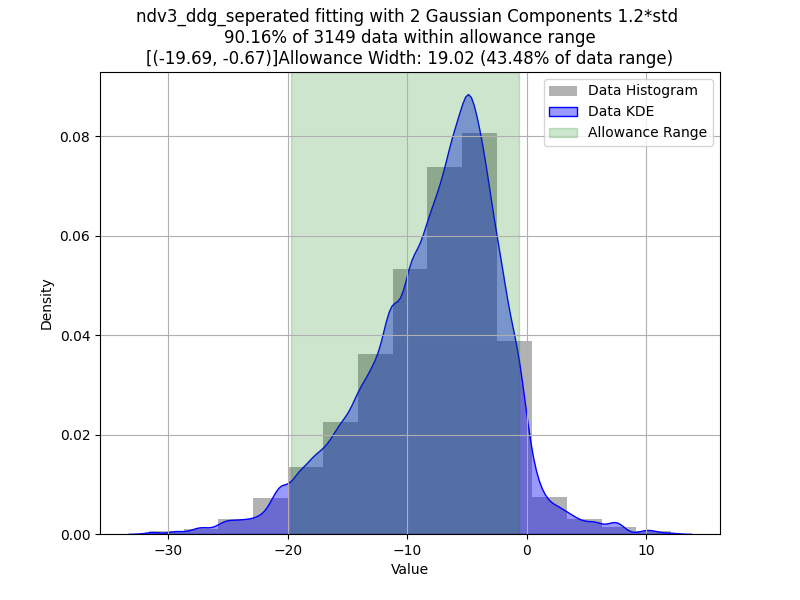

Negative samples were generated by computational design. A Rosetta design protocol named *new residue design* is used to generate sugar binding proteins that include CH-π interactions. In stage 1, it finds a suitable pocket for a ligand in a given protein scaffold and alters the interacting residue to alanine. Then, it designs a protein sequence to have proper polar and CH-π interactions to target sugar in stage 2. Selecting the top 10% of the worst designed proteins in both stage 1 and stage 2 according to Rosetta binding free energy. Those would be the negative binder database, with either lack of interaction or having inappropriate interaction to sugar.

### Supplementary Data 5. Model architecture

Supplementary figure 2 The architecture of CLIMBS classifier model

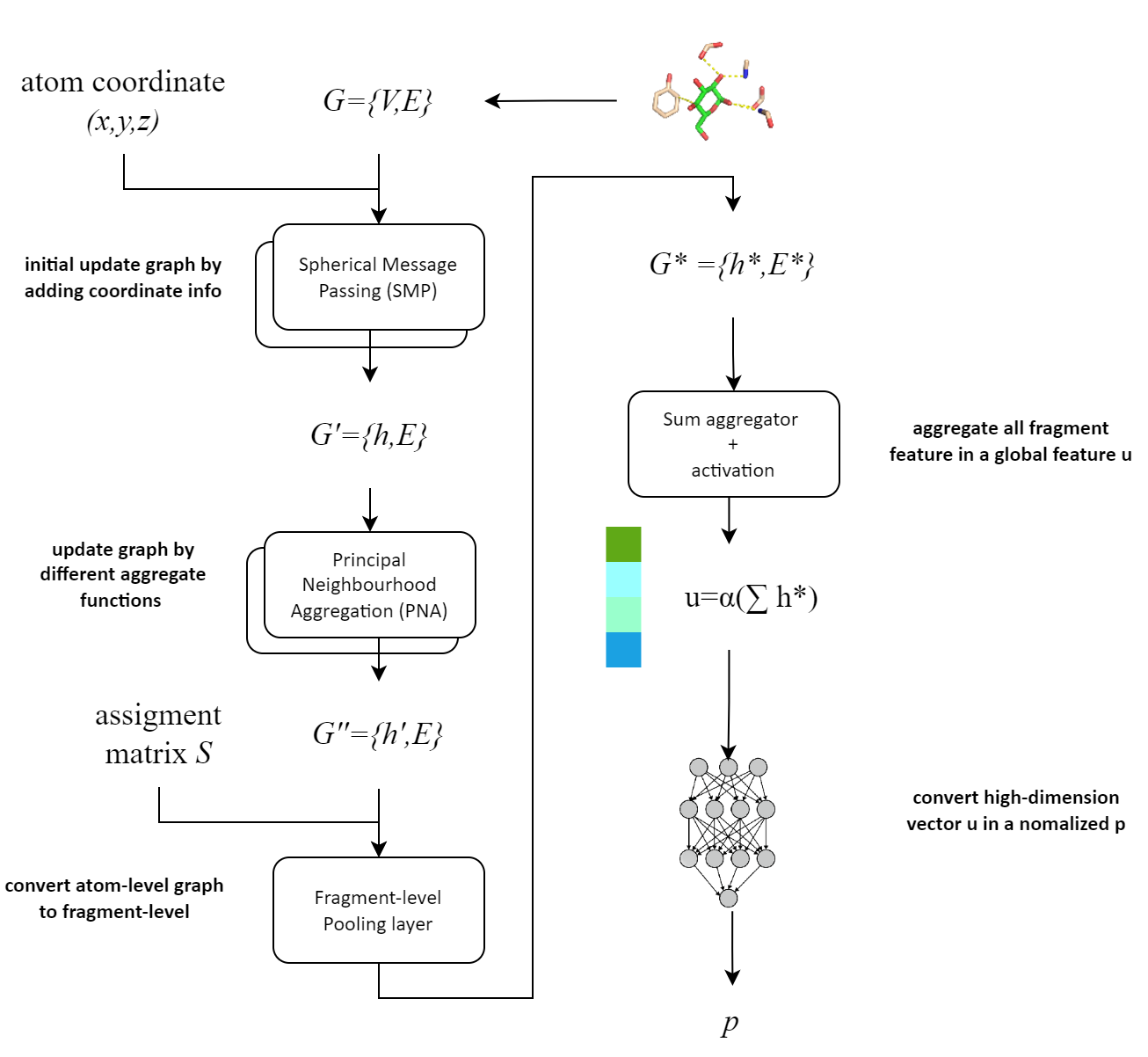

The minimum binding site information is inputted to the model including atom coordinates of structure, an interaction graph G, and an atom-to-fragment assignment matrix S. After several blocks (SMP, PNA, pooling layers and sum aggregator), the classifier model outputs a normalized p. The activation function (α) is rectified linear unit (ReLU) in the model.

### Supplementary Data 6. Pooling layer selection

As a result in Supplementary Table 3, all three models have similar performance for mono-saccharide samples prediction, with the highest accuracy of 99.52% for the fragment-level model and the lowest accuracy of 96.38% for the molecule-level model. For di-saccharide samples prediction, fragment-level model is apparently better than the other, with the accuracy of 81.25% that 6.25% and 12.50% higher than molecule-level and atom-level respectively.

Supplementary table 3 Accuracy and training loss of CLIMBS with different pooling layers. Classifier here were trained on GlcNAc, GlcA and GlcNAc-GlcA (db_p2, Supplementary Data 3b). The 2^nd^ and 3^rd^ columns are the testing results on GlcNAc and GlcA, the 4^th^ and 5th columns are on GlcNAc-GlcA di-saccharides**.**

| Pooling layer | mono-sac loss | mono-sac accuracy | di-sac loss | di-sac accuracy |
| --- | --- | --- | --- | --- |
| atom-level | 0.022 | 98.07% | 0.277 | 68.75% |
| fragment-level | 0.019 | 99.52% | 0.19 | 81.25% |
| molecule-level | 0.056 | 96.38% | 0.245 | 75.00% |

### Supplementary Data 7. Robustness check

From the pooling layer selection assay, we knew CLIMBS performed quite well on trained sugar type samples. To make the model more fragile and sensitive during evaluation, a model that trained on GlcNAc and GlcA was used to evaluate glucose (Glc) (for detail see Supplementary Data 3c). The model had an initial accuracy of 76.36% (Supplementary table 4), but different disturbances were introduced to the test set. Missing the CH-π interacting information has almost no effect, but missing all interaction information or removing CH-π interacting residue decreases 4% of the accuracy, with a drop of true positive predictions. Moreover, adding an additional CH-π interacting residue significantly decreases the amount of true negatives with an accuracy drop of -19.17%. Although the disturbances of CH-π residue harm the CLIMBS performance, the accuracies of attacked samples (72.00% and 57.19%) are still higher than 50%.

Supplementary table 5 Accuracy and training loss of CLIMBS facing adversarial attacks. Model here was trained on GlcNAc and GlcA, and tested by Glc samples with missing or addition information (db_p2, see Supplementary Data 3c). TP: true positive, FN: false negative, TN: true negative, FP: false positive.

| **Disturbance** | **Loss** | **Accuracy** | **TP** | **FN** | **TN** | **FP** |
| --- | --- | --- | --- | --- | --- | --- |
| -- | 0.297 | 76.36% | 403 | 57 | 298 | 162 |
| remove CH-π interactions | 0.299 | 76.47% | 404 | 56 | 298 | 162 |
| remove all  interactions | 0.319 | 72.22% | 368 | 92 | 295 | 165 |
| remove CH-π residues in positive sample | 0.315 | 72.00% | 363 | 97 | 298 | 162 |
| add CH-π residues in negative sample | 0.429 | 57.19% | 403 | 57 | 122 | 338 |
|  |  |  | 460 | | 460 | |

### Supplementary Data 8. Calibrating other methods

Every method needs to set a threshold value to separate predicted binding and unbound, since Rosetta Energy Function(*8*), Autodock4 score function(*9*), HADDOCK score function(*10*) and DIFFDOCK(*11*) confidence model return a scalar value rather than a binary value.

We investigated the effect of threshold value on model accuracy, sensitivity, specificity, precision and F1 score of all methods. To get the best performance of each model, the threshold value that has the maximum accuracy was chosen (supplementary table 2 and supplementary figure 1).

Supplementary table 2 Threshold and normalised threshold value of each method to predict as bound.

|  | Threshold | Normalised threshold |
| --- | --- | --- |
| CLIMBS | >0.0159 | >0.0159 |
| Rosetta binding free energy | <-12.9889 kcal/mol | <0.597 |
| Autodock4 binding free energy | <-3.5982 kcal/mol | <0.7188 |
| HADDOCK3 binding free energy | <-26.2725 kcal/mol | <0.4908 |
| DIFFDOCK confidence | <-3.3255 | <0.3008 |

Supplementary figure 3 Normalised score calibration for positive sample and negative sample for CLIMBS and other methods.

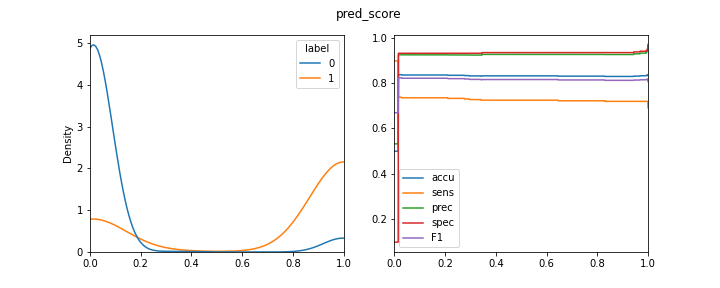

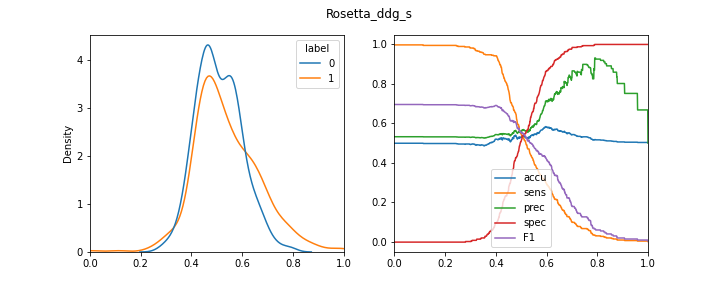

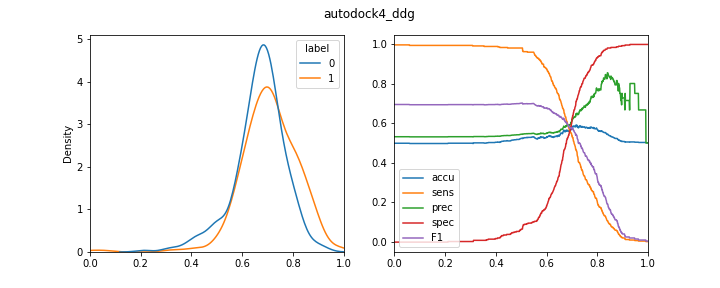

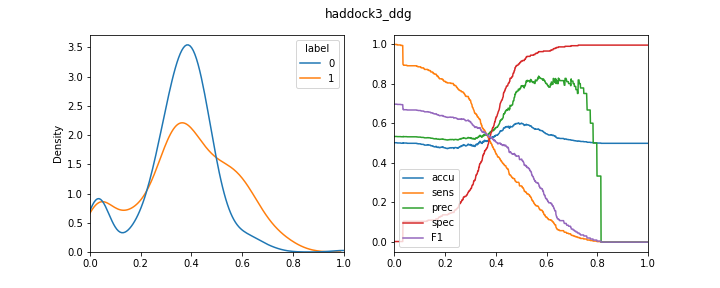

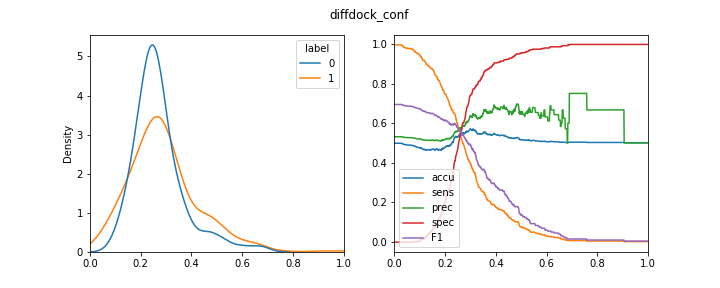

Scores from each method were normalised between 0 and 1, corresponding to the lowest and highest scoring samples. Distributions of normalised score plot as kernel density estimate. Positive sample shows in orange and negative sample shows in blue. (right) Relationship between normalized threshold value and model indicators. accu: accuracy, sens: sensitivity, spec: specificity, prec: precision, F1: F1 score. pred_score: sugar binding classifier score, diffdock_conf: DIFFDOCK confidence, Rosetta_ddg: Rosetta binding free energy, autodock4_ddg: Autodock4 binding free energy, haddock3_ddg: HADDOCK3 binding free energy.

### Supplementary Data 9. CLIMBS on docking and design problems

The carbohydrate-protein complexes were generated by Chai-1 (*12*). Proteins amino acid sequence and carbohydrates SMILES sequence were input without MSA, template and restraints. For docking, rank 0 model of each carbohydrate-protein complex with protein pLDDT>90, ligand pLDDT>70, protein backbone RMSD <2 Å to reference structure was kept. For design, rank 0 model of each carbohydrate-protein complex with protein pLDDT>90, ligand pLDDT>70 was kept. Evaluation by metrics was performed on Rosetta relaxed structure.

CLIMBS performance was tested on the *db_dock* dataset and compared with Rosetta ddg at different energy cutoff. CLIMBS_D was a model trained on a selected number of complexes with resolution < 2 Å (*db_w2*).

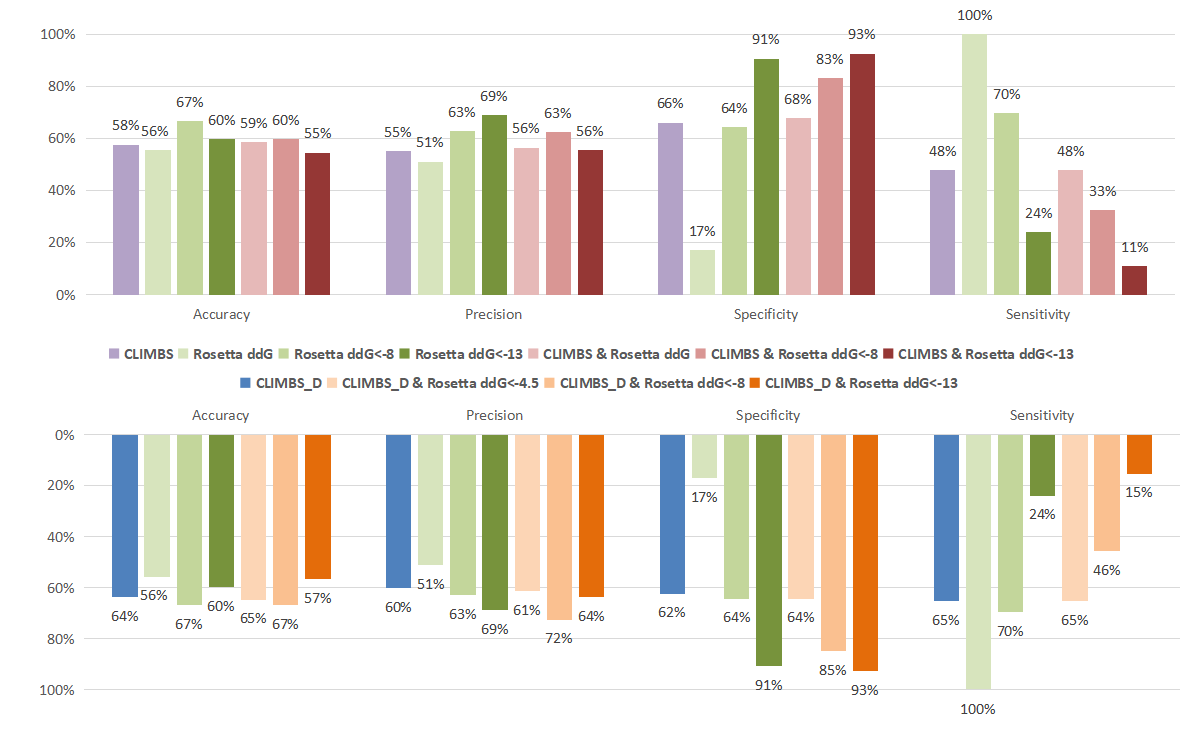

Supplementary figure 4 Performance of CLIMBS, CLIMBS_D and Rosetta on carbohydrate-protein docking problem. CLIMBS_D was trained by less samples withs high structure quality (db_w2, see Supplementary Data 3l) . 99 samples were included. Structures with ligand RMSD <2 Å after superimposing the protein to the referral structure are regard as bound. Thresholds to predict as positive in for each metric: CLIMBS=1, CLIMBS_D=1, Rosetta score function ddG <-4.5, <-8 and <-13. The positive predictions of combined metrics are the AND subset of each positive predictions. accuracy: (TP+TN)/(TP+FP+TN+FN), precision: TP/(TP+FP), specificity: TN/(TN+FP), sensitivity: TP/(TP+FN). TP: True Positive, TN: True Negative, FP: False Positive, FN: False Negative.

Supplementary table 6 Label of carbohydrate-protein complexes in design problem.1: bound, 0: unbound. Previous research (13) did a cross-binding assay between different carbohydrate and 4 protein variants. Carbohydrate-protein results with y-median fluorescence intensity higher than 5000 were regarded as bound.

|  | 08F | 13D | 24i | 83H |
| --- | --- | --- | --- | --- |
| Chitobiose | 1 | 1 | 1 | 0 |
| Core5 | 0 | 1 | 1 | 0 |
| Core8 | 0 | 0 | 1 | 0 |
| GAL | 0 | 0 | 0 | 0 |
| GalNAc | 0 | 0 | 1 | 0 |
| GlcNAc | 0 | 0 | 0 | 0 |
| H3 | 0 | 0 | 1 | 0 |
| LacNAc | 0 | 0 | 0 | 0 |
| Lec | 1 | 1 | 1 | 0 |
| Man | 0 | 0 | 0 | 0 |
| Neu5Ac | 0 | 0 | 0 | 0 |
| Neu5Gc | 0 | 0 | 0 | 0 |
| TF | 1 | 1 | 0 | 0 |

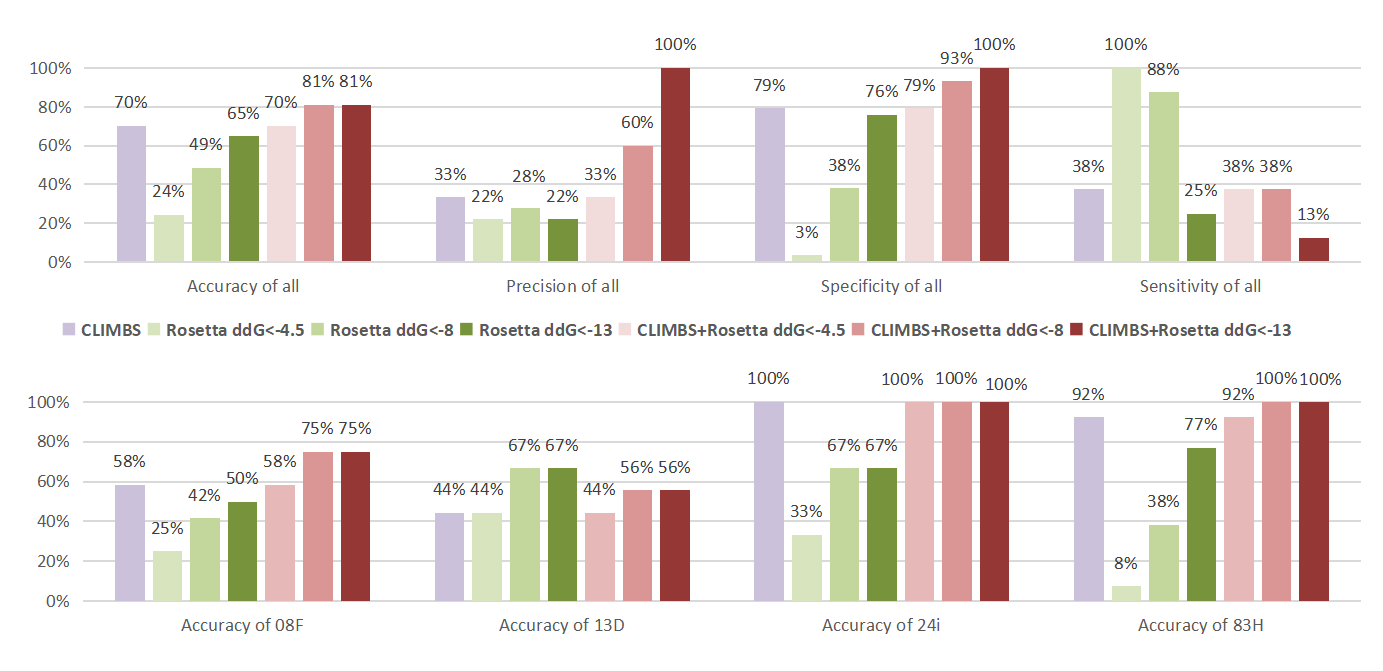

Supplementary figure 5 Performance of different scoring methods on design carbohydrate-binding protein problem. In total 37 samples were included: 12 samples of 08F, 9 samples of 13D, 3 samples of 24i, 13 samples of 83H. Four protein variants bind to different carbohydrate in 13 candidates. Thresholds to predict as positive in for each metric: CLIMBS=1, Rosetta score function ddG <-4.5, <-8 and <-13. The positive predictions of combined metrics are the AND subset of each positive predictions. accuracy: (TP+TN)/(TP+FP+TN+FN), precision: TP/(TP+FP), specificity: TN/(TN+FP), sensitivity: TP/(TP+FN). TP: True Positive, TN: True Negative, FP: False Positive, FN: False Negative

### Supplementary method 1. Rosetta relaxation and docking(14)

a. Rosetta relaxation command

*rosetta_scripts.linuxgccrelease -nstruct 1 -parser:protocol single_sugar_relax_fast.xml -include_current -ignore_unrecognized_res -include_sugars -s file_name*

b. Rosetta relaxation *script single_sugar_relax_fast.xml*

*<ROSETTASCRIPTS>*

*<RESIDUE_SELECTORS>*

*<Chain name='sugar' chains='A' />*

*<Not name="protein" selector="sugar"/>*

*<InterfaceByVector name="strict_interface" grp1_selector="sugar" grp2_selector="protein"/>*

*<PrimarySequenceNeighborhood name="interface" selector="strict_interface"/>*

*</RESIDUE_SELECTORS>*

*<MOVERS>*

*<EnsureExclusivelySharedJumpMover name="set_sugar_other_jump" residue_selector="sugar" />*

*<AtomTree name="set_foldtree"	docking_ft="1" />*

*<InterfaceAnalyzerMover name='interface_analyis' ligandchain='A' interface_sc='1' tracer='0'/>*

*</MOVERS>*

*<FILTERS>*

*<Ddg name="ddg" confidence="0" repeats='1' />*

*</FILTERS>*

*<PROTOCOLS>*

*<Add mover_name="set_foldtree"/>*

*<Add mover_name="set_sugar_other_jump"/>*

*<Add mover="relax"/>*

*<Add mover="interface_analyis" />*

*<Add filter="ddg"/>*

*</PROTOCOLS>*

*<OUTPUT />*

</ROSETTASCRIPTS>

c. Rosetta docking command

*rosetta_scripts.linuxgccrelease -s file_name -in:file:native file_name -parser:protocol local_ligand_dock.xml -nstruct 200 -include_sugars -ex1 -ex2*

d. Rosetta docking script *local_ligand_dock.xml*

*<ROSETTASCRIPTS>*

*<SCOREFXNS>*

*<ScoreFunction name="ligand_soft_rep" weights="ligand_soft_rep">*

*<Reweight scoretype="fa_elec" weight="0.42"/>*

*<Reweight scoretype="hbond_bb_sc" weight="1.3"/>*

*<Reweight scoretype="hbond_sc" weight="1.3"/>*

*<Reweight scoretype="rama" weight="0.2"/>*

*</ScoreFunction >*

*<ScoreFunction name="hard_rep" weights="ligand">*

*<Reweight scoretype="fa_intra_rep" weight="0.004"/>*

*<Reweight scoretype="fa_elec" weight="0.42"/>*

*<Reweight scoretype="hbond_bb_sc" weight="1.3"/>*

*<Reweight scoretype="hbond_sc" weight="1.3"/>*

*<Reweight scoretype="rama" weight="0.2"/>*

*</ScoreFunction>*

*</SCOREFXNS>*

*<LIGAND_AREAS>*

*<LigandArea name="docking_sidechain" chain="X" cutoff="6.0" add_nbr_radius="true" all_atom_mode="true" minimize_ligand="10"/>*

*<LigandArea name="final_sidechain" chain="X" cutoff="6.0" add_nbr_radius="true" all_atom_mode="true"/>*

*<LigandArea name="final_backbone" chain="X" cutoff="7.0" add_nbr_radius="false" all_atom_mode="true" Calpha_restraints="0.3"/>*

*</LIGAND_AREAS>*

*<INTERFACE_BUILDERS>*

*<InterfaceBuilder name="side_chain_for_docking" ligand_areas="docking_sidechain"/>*

*<InterfaceBuilder name="side_chain_for_final" ligand_areas="final_sidechain"/>*

*<InterfaceBuilder name="backbone" ligand_areas="final_backbone" extension_window="3"/>*

*</INTERFACE_BUILDERS>*

*<MOVEMAP_BUILDERS>*

*<MoveMapBuilder name="docking" sc_interface="side_chain_for_docking" minimize_water="true"/>*

*<MoveMapBuilder name="final" sc_interface="side_chain_for_final" bb_interface="backbone" minimize_water="true"/>*

*</MOVEMAP_BUILDERS>*

*<RESIDUE_SELECTORS>*

*<InterfaceByVector name="interface">*

*<Chain chains="X"/>*

*<Chain chains="B"/>*

*</InterfaceByVector>*

*</RESIDUE_SELECTORS>*

*<FILTERS>*

*<Ddg name="ddg" confidence="0"/>*

*<Sasa name="SASA" confidence="0"/>*

*<ResidueCount name="resi_interface" residue_selector="interface" confidence="0"/>*

*</FILTERS>*

*<MOVERS>*

*<Translate name="translate" chain="X" distribution="uniform" angstroms="5.0" cycles="50" force="true"/>*

*<Rotate name="rotate" chain="X" distribution="uniform" degrees="360" cycles="500"/>*

*<SlideTogether name="slide_together" chains="X"/>*

*<HighResDocker name="high_res_docker" cycles="1" repack_every_Nth="1" scorefxn="ligand_soft_rep" movemap_builder="docking"/>*

*<FinalMinimizer name="final" scorefxn="hard_rep" movemap_builder="final"/>*

*<InterfaceScoreCalculator name="add_scores" chains="X" scorefxn="hard_rep"/>*

*<AddJobPairData name="system_name" key="system_name" value_type="string" value_from_ligand_chain="X" />*

*<ParsedProtocol name="low_res_dock">*

*<Add mover_name="translate"/>*

*<Add mover_name="rotate"/>*

*<Add mover_name="slide_together"/>*

*</ParsedProtocol>*

*<ParsedProtocol name="high_res_dock">*

*<Add mover_name="high_res_docker"/>*

*<Add mover_name="final"/>*

*</ParsedProtocol>*

*<ParsedProtocol name="reporting">*

*<Add mover_name="add_scores"/>*

*</ParsedProtocol>*

*</MOVERS>*

*<PROTOCOLS>*

*<Add mover_name="high_res_dock"/>*

*<Add mover_name="reporting"/>*

*<Add filter="ddg" />*

*<Add filter="SASA" />*

*<Add filter="resi_interface" />*

*</PROTOCOLS>*

*</ROSETTASCRIPTS>*
